# Electrostatic-driven Interactions Enhance Intratumoral Retention and Antitumor Efficacy of Immune Checkpoint Blockade Antibodies

**DOI:** 10.1101/2023.12.22.573144

**Authors:** Rashmi P Mohanty, Yuting Pan, Mae M Lewis, Melissa Soto, Esther Y Maier, Riyad F Alzhrani, Debadyuti Ghosh

**Affiliations:** Division of Molecular Pharmaceutics and Drug Delivery, College of Pharmacy, The University of Texas at Austin; Austin, Texas, USA; Department of Biomedical Engineering, The University of Texas at Austin; Austin, Texas, USA; Drug Dynamics Institute, The University of Texas at Austin; Austin, Texas, USA

## Abstract

Tumor extracellular matrix (ECM) forms a net negative charged network that interacts with and hinders the transport of molecules partly based on electrostatic interactions. The focus on drug delivery in solid tumors has traditionally been on developing neutral charge coatings to minimize interactions with the ECM for improved transport. In contrast to this prior work, we recently found a cationic peptide that interacted electrostatically with the negatively charged components of the ECM, resulting in enhanced uptake and retention of nanoparticles in tumor ECM and tumor tissue. Based on this previous study, here, we hypothesize that the electrostatically driven interactions of the cationic peptide will improve the binding and retention of immune checkpoint blockade antibodies (ICBs), ultimately enhancing their antitumor immunogenic responses. We prepared peptide antibody (Ab) conjugates by conjugating the cationic peptide to ICBs, anti-cytotoxic T lymphocyte antigen 4 (∝-CTLA4) and anti-programmed cell death ligand-1 (∝-PD-L1) Abs, using copper-free click chemistry. We confirmed an average of 1 – 2 peptides per Ab. The cationic peptide electrostatically interacted with the net negatively charged tumor ECM and improved the binding of the Abs to the tumor ECM without affecting their antigen recognition capacities. Modifying the Abs due to cationic peptide conjugation reduced the systemic exposure of the Abs and did not induce treatment-related toxicities. We quantified a significantly higher population of tumor-infiltrating CD8^+^ T cells and a significant depletion of regulatory T cells in the tumor and tumor-draining lymph nodes upon peptide conjugation, which resulted in a better therapeutic outcome of the ICBs.

**ONE SENTENCE SUMMARY:** Electrostatic interaction-based intratumoral retention enhances antitumor responses of immune checkpoint blockade antibodies upon local administration.

## INTRODUCTION

Immune checkpoint blockade (ICB) has been transformative in cancer therapy to achieve long-term, durable remissions in patients with tumors such as melanoma. During homeostasis, immune checkpoints are inhibitory pathways of the immune system to maintain self-tolerance and protect host tissues from collateral damage during anti-microbial immune responses. In cancers, this mechanism is unfortunately exploited to evade immune-mediated rejections, leading to their progression.^1^ Antibodies (Abs) that inhibit immune checkpoints, such as ∝-CTLA4 and ∝-PD-L1/anti-programmed cell death-1 (∝-PD-1), can unleash antitumor immunity and have been “breakthrough” therapies approved for clinical use for the treatment of various types of cancer, including melanoma.^2–7^ The first successful immunotherapy to block the negative regulatory pathway on T cells is the ∝-CTLA4 Ab, which was approved by the FDA to treat metastatic melanoma.^8^ CTLA4 is an activation-induced T cell surface molecule with homology to CD28 and is expressed on activated CD4^+^ and CD8^+^ T cells and regulatory T cells (T_regs_)^9^ as an inducer of immunosuppressive signals.^10^ Blocking CTLA4 using antibodies (∝-CTLA4) can potentiate the activation of effector T cells (T_eff_) and deplete T_regs_ for antitumor efficacy.^9^ Additional success was reported with immune checkpoint blockade with antibodies against PD-1 and PD-L1.^6, 11^ Here, tumor cells and antigen-presenting cells can express PD-L1, which bind to PD-1 on activated T cells to evade immune response.^12, 13^ The PD-L1/PD-1 interaction decreases cytokine production, inhibits proliferation of T cell receptor-mediated lymphocyte proliferation, and induces apoptosis of T cells.^12, 14^ Both ∝-PD-1 and ∝-PD-L1 blocking Abs have been developed to inhibit tumor growth in mouse models and in the clinic.^13^ Also, since PD-L1/PD-1 and CTLA4 checkpoints are non-redundant, the combination of ∝ −CTLA4 and ∝ −PD-L1/∝ −PD-1 can have synergistic antitumor effects, thereby shifting the tumor microenvironment from suppressive to inflammatory.^2, 3, 5, 15^

While ICB is promising, systemic delivery of immune checkpoint inhibitors and other immunotherapies often leads to undesired toxicity and immune-related adverse events, which limit their efficacy in the clinic. For example, in a phase II clinical trial, interleukin-12 administered systemically triggered cytokine release syndrome, which resulted in vascular leakage and led to the failure of the trial ^16^. In another clinical trial, a 10 mg/kg dose of ∝-CTLA4 ipilimumab given systemically resulted in greater severity of immune-related adverse events experienced by melanoma patients than a lower dose due to increased systemic exposure into healthy tissues.^17, 18^ Combination (∝-PD-1 and ∝-CTLA4) therapy can be given to those who do not respond to monotherapy, but the combination regimen even further increases the severity of immune-related adverse events and toxicities.^3, 19, 20^ In the CHECKMATE 068 trial, 59% of patients reported severe treatment-related adverse events, such as increased grade 3-4 alanine aminotransferase and diarrhea, with systemic combination ∝-PD-1 nivolumab and ∝-CTLA4 ipilimumab therapy compared to 23% and 28% with ∝-PD-1 and ∝-CTLA4 monotherapy, respectively.^20^

Local delivery of ICBs via intratumoral,^21–24^ peritumoral^25, 26^ and subcutaneous^27^ administration has improved antitumor efficacy compared to systemic delivery due to reduced systemic exposure and ability for increased dosing. Also, there are preclinical and clinical evidence that intratumoral immunomodulation through local anchoring of antibodies to the tumor microenvironment can generate systemic antitumor immune response.^28^ Ishihara et al. conjugated a collagen-binding peptide to ICBs (∝-CTLA4 and ∝-PD-L1 Abs) to enhance their retention in the tumor extracellular matrix (ECM).^29^ The enhanced retention of the Abs in tumor tissue enhanced the antitumor efficacy of the ICBs in melanoma and breast tumors by improving T cell activation and reducing the side effects of therapy. Additionally, they reported that via local administration, there was systemic antitumor immunity, which has the potential for the treatment of metastatic tumors.^29^

We recently demonstrated that a cationic peptide (CRRRRKSAC, net charge +5) enabled enhanced diffusion, uptake, and retention of a phage (i.e., bacterial virus) in tumor ECM and *ex vivo* tumor tissue compared to a neutral peptide (CKPGDGGPC, net charge 0).^30^ The cationic peptide interacted electrostatically with the net negatively charged tumor ECM, resulting in a steeper concentration gradient than with neutral charged counterparts for greater penetration while achieving improved retention in tumor tissue. Similar phenomena have previously been reported in other highly negatively charged biological barriers, such as cartilage^31, 32^ and mucus.^33^ Weak and reversible interaction with the surrounding environment provides higher retention and deeper tissue penetration by taking advantage of the electrostatic interactions.^34, 35^

Building from this observation, we hypothesized the cationic peptide could be used as a carrier to improve the retention of ICBs in the tumor ECM. In this study, we aimed to use the cationic peptide discovered in our previous work to enhance the delivery and therapeutic outcome of ∝-CTLA4 and ∝-PD-L1 Abs in a murine melanoma model. The cationic peptide was expected to interact weakly with the tumor ECM electrostatically, thereby enhancing the binding of the ICBs to the ECM. The enhanced binding helped with the retention of ICBs in tumor tissue upon local administration and reduced their systemic exposure compared to free, unconjugated antibodies, which is critical to avoid toxicity. Finally, we used cationic peptide conjugates to improve immune profiling in tumor microenvironment and antitumor efficacy.

## RESULTS

### Conjugation of peptides to ICBs

We conjugated the cationic and neutral peptides to the free amines of the ICBs using two-step copper-free click chemistry (**Fig. 1a**). We calculated the concentration of unmodified and peptide conjugated ICBs from their absorbance measurements at 280 nm. Conjugation of the peptides to the Abs was confirmed from reduced SDS-PAGE with subsequent determination of their molecular weight from LC-MS. From the bands on SDS-PAGE, the slight increase in both the light chain (25 kDa) and the heavy chain (50 kDa) of the ICBs indicated peptide conjugation to both heavy and light chains of the Abs (**Fig. 1b**). The molecular weight of the cationic and the neutral peptides was 1287.55 Da and 985.12 Da, respectively. The increase in the molecular weight of the Abs was confirmed using LC-MS (**Fig. 1c**). The Abs were reduced before performing LC-MS to observe noticeable changes in the molecular weight of both light and the heavy chain upon peptide conjugation. Upon conjugation of the cationic peptide, the molecular weight of the heavy chain of ∝-CTLA4 Ab increased from 51737 Da to 52990 Da, and the light chain increased from 23978 Da to 25532 Da. Similarly, upon conjugation of the neutral peptide, the molecular weight of the heavy chain of ∝-CTLA4 Ab increased to 52988 Da, and the light chain increased to 25231 Da. Based on the shift in molecular weight from peptide conjugation, an average of 2.2 cationic peptides and 2.6 neutral peptides were conjugated to ∝-CTLA4 Ab. The molecular weight of the heavy chain of the ∝-PD-L1 Ab increased from 50311 Da to 51736 Da upon conjugation of cationic peptide, and the molecular weight of the light chain increased from 23618 Da to 24869 Da upon conjugation of neutral peptide. On average, 1.1 cationic peptides and 1.3 neutral peptides were conjugated to the ∝-PD-L1 Ab.

**Fig. 1.**
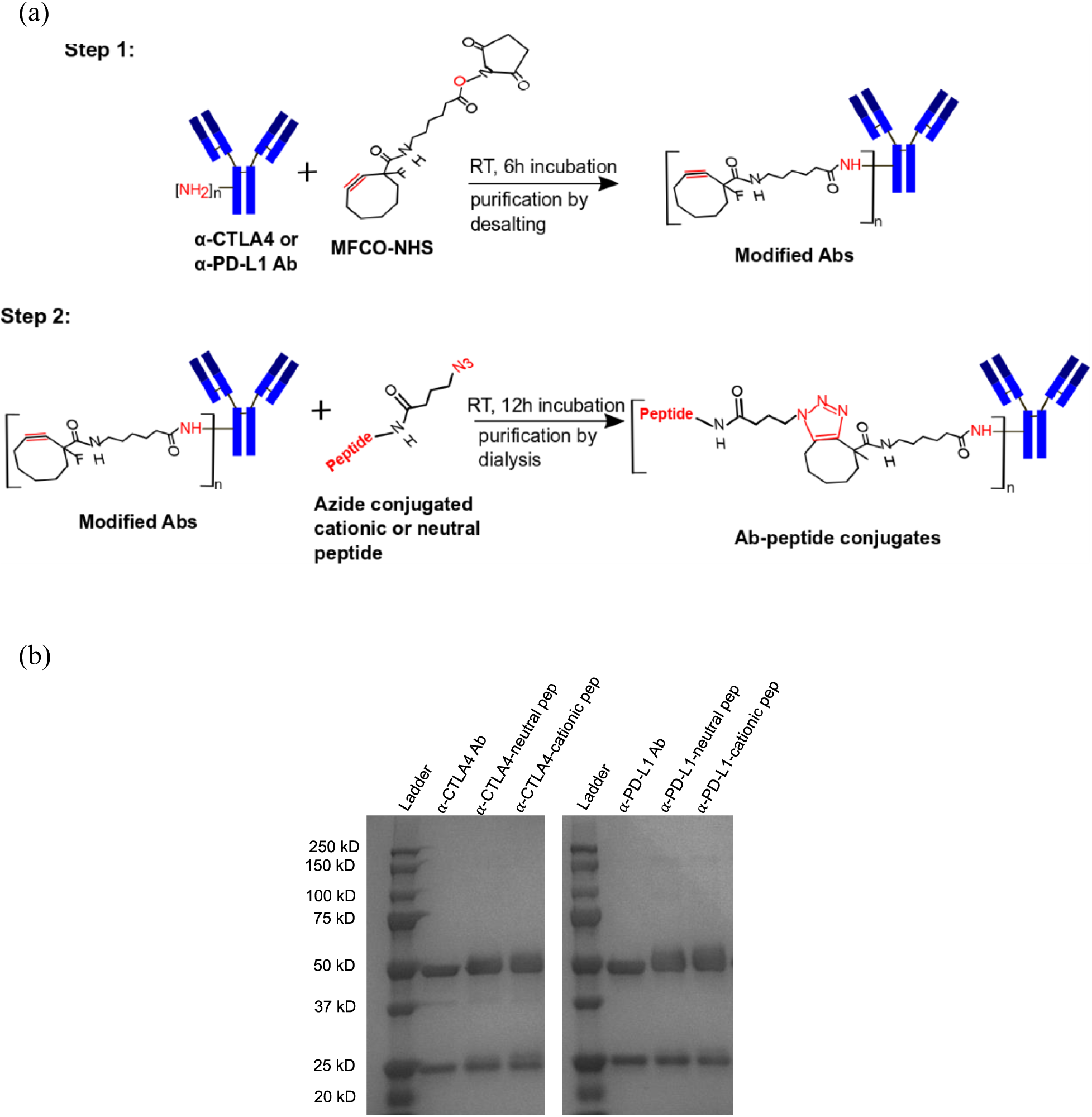
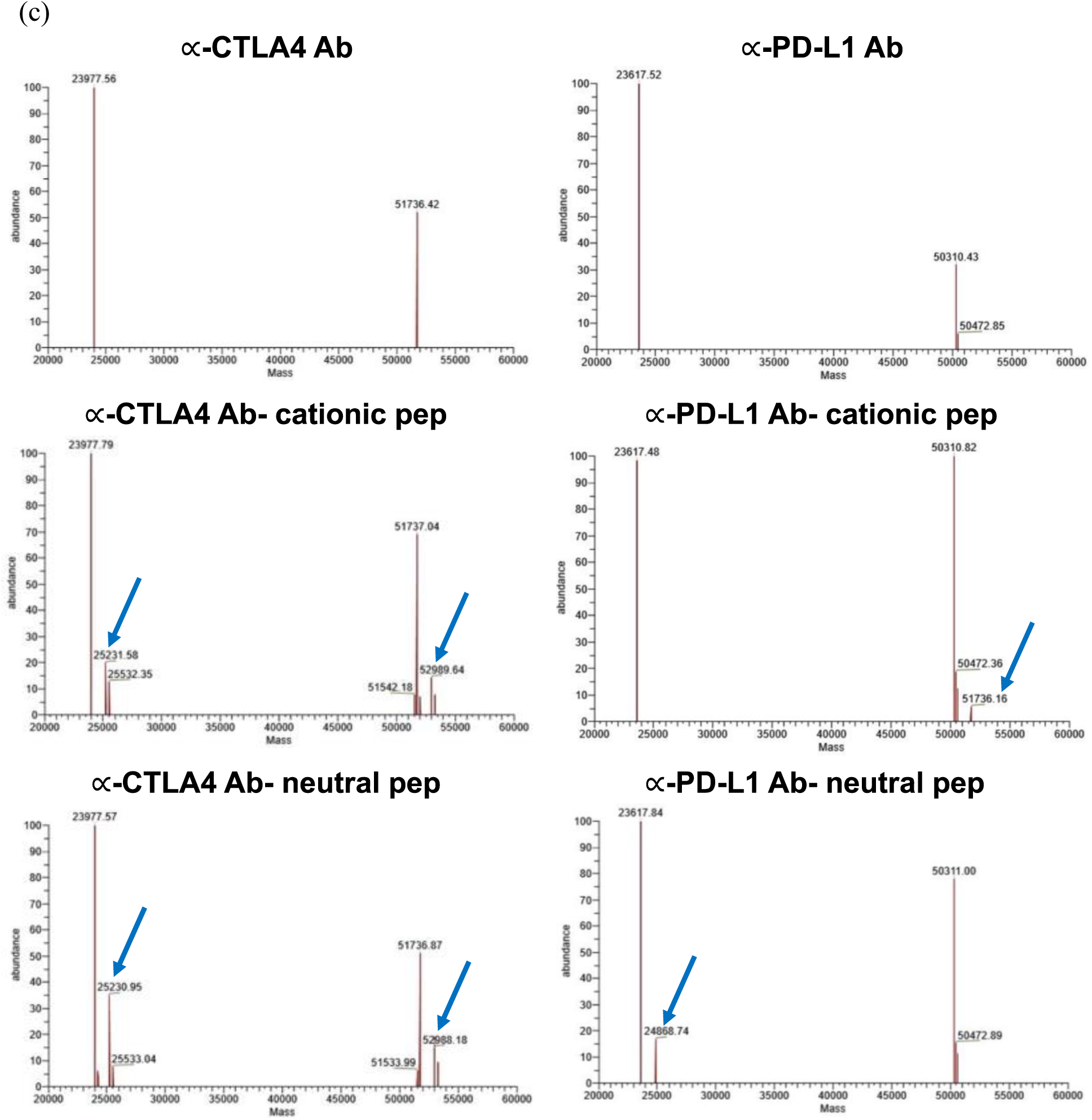
On an average, 1-2 peptides were conjugated to the ICBs using copper-free click chemistry. **(a)** Antibody (Ab)-peptide conjugates were synthesized using a two-step copper-free click chemistry process. In step 1, ICBs ∝-CTLA4 and ∝-PD-L1 were modified by incubating with MFCO-NHS for 6 h at RT while shaking at 2 rpm. The modified Abs were purified by gel filtration using a PD-10 desalting column. In step 2, the modified Abs were reacted with either azide modified cationic or neutral peptides. The reaction was carried out at RT for 12 h while gently shaking at 2 rpm. After, the Ab-peptide conjugates were purified using dialysis. Peptide conjugation to Abs was verified using reduced SDS-PAGE and LC-MS. **(b)** Image of the reduced SDS-PAGE confirmed an increase in molecular weight of both light (25 kDa) and heavy (50 kDa) chains of the Abs after peptide conjugation. The observed slight increase in the molecular weight of Abs is from the molecular weight (∼1 kDa) of the peptides. **(c)** From LC-MS analysis, it was confirmed that on average, approximately two cationic and neutral peptides were conjugated to the ∝-CTLA4 Ab; only one cationic peptide was conjugated to the heavy chain of the ∝-PD-L1 Ab; and one neutral peptide was conjugated to the light chain of the ∝-PD-L1 Ab.

### Cationic peptide enhanced binding of ICBs to the tumor-like ECM without compromising the functionality of the ICBs

It is important that upon peptide conjugation to ICBs, there is no loss of antibody binding to its cognate antigen. To confirm this, we measured and compared the binding affinities of unconjugated and peptide-conjugated ICBs to their target antigens and to tumor ECM by ELISA (**Fig. 2**). The binding affinity (k_d_) of ∝-CTLA4 Ab with its target antigen rmCTLA-4 is 361 pM. Upon conjugation of either cationic or neutral peptide, the k_d_ was 302 pM and 590 pM, respectively (**Fig. 2a**). Similarly, k_d_ of ∝-PD-L1 Ab to its target antigen rmPD-L1 was 53 pM. Upon conjugation of either cationic or neutral peptide to ∝-PD-L1 Ab, the k_d_ to the target antigen was 85 pM and 102 pM, respectively (**Fig. 2b**). The similar values of the binding affinities suggest that the peptide conjugation did not impair antigen binding of the Abs.

**Fig. 2.**
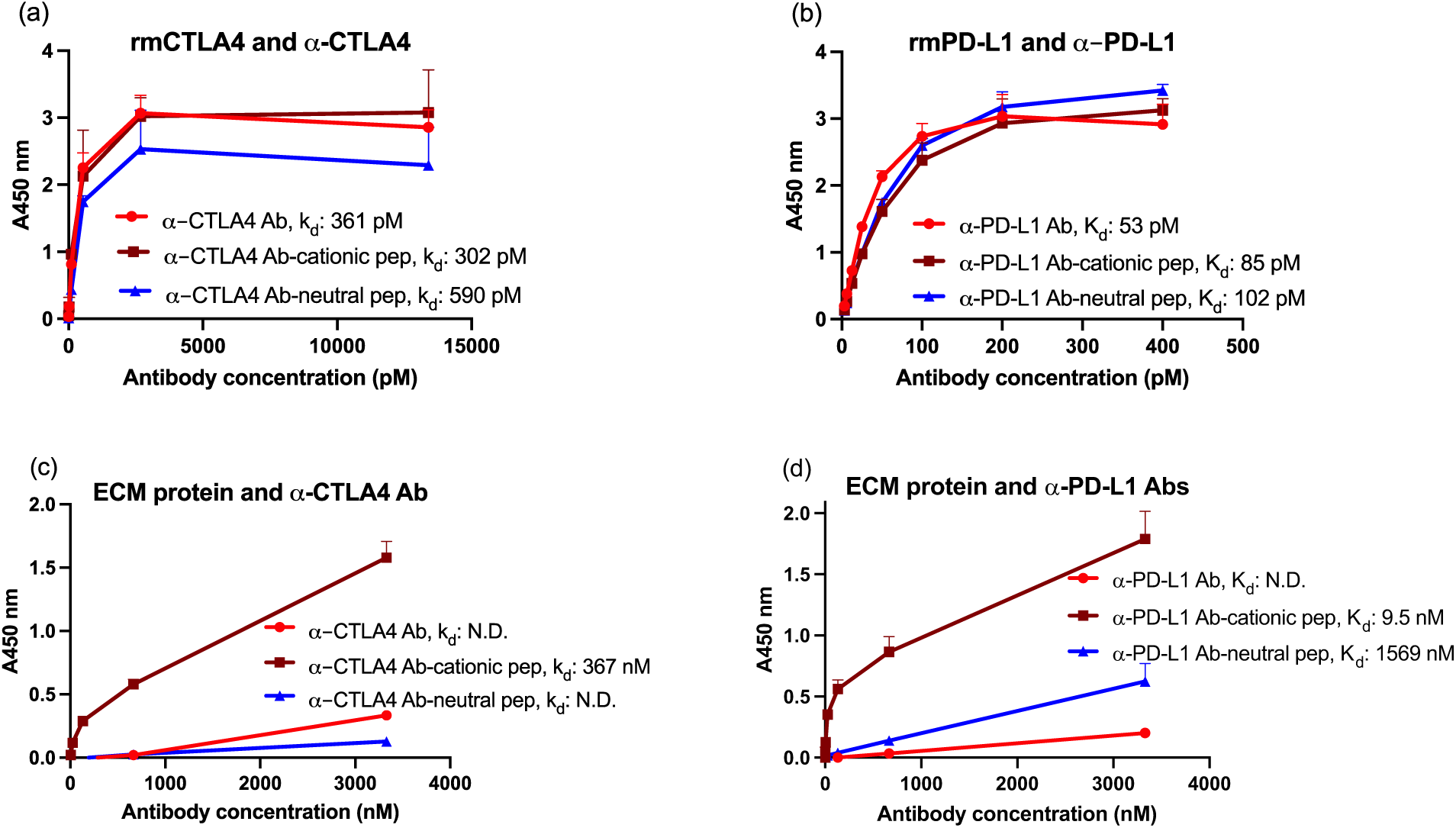
Cationic peptide conjugation enhanced the binding of ICBs to the ECM while preserving the antigen recognition ability of ICBs. Binding affinities of unconjugated and peptide-conjugated ICBs were quantified using enzyme-linked immunosorbent assay (ELISA). Absorbance at 450 nm (A450 nm) for binding of (**a**) unconjugated and peptide conjugated ∝-CTLA4 Ab with its target antigen rmCTLA4, (**b**) unconjugated and peptide conjugated ∝-PD-L1 Ab with its target antigen rmPD-L1, (**c**) unconjugated and peptide conjugated ∝-CTLA4 Ab with ECM protein, and (**d**) unconjugated and peptide conjugated ∝-PD-L1 Ab with ECM protein was plotted with Ab concentration. Binding affinities (k_d_) were estimated based on non-linear least square regression analysis. N.D. is not determined because of the low signal. The data were presented as mean±SD (n=4).

We then confirmed that the peptide-conjugated Abs can bind to the net-negatively charged tumor-like ECM. The binding affinities of the free and peptide-conjugated Abs were measured and calculated against a tumor-like ECM model. The tumor ECM was consisted of collagen and hyaluronic acid (HA) at a concentration of 10 µg/mL and 1.05 µg/mL, respectively. Collagen and HA are the two most abundant components of tumor ECM that restrict drug delivery.^36–39^ The unconjugated, free Abs did not bind to the tumor ECM, which was reflected by the smaller absorbance values observed throughout the studied Ab concentration range (**Figs. 2c and 2d**); there was no k_d_ that could be calculated. Upon conjugation of the neutral peptide to Abs, neutral peptide-conjugated ∝-CTLA4 Ab did not bind to the tumor ECM, and the neutral peptide-conjugated ∝-PD-L1 Ab bound weakly to the tumor ECM with k_d_ of 1569 nM. Interestingly, upon conjugation of cationic peptide, both ∝-CTLA4 Ab and ∝-PD-L1 Ab bound to tumor ECM; the k_d_ decreased to 367 nM and 9.5 nM, respectively. The results from the binding assays demonstrate that peptide conjugation does not adversely affect antigen binding of ICBs, which suggests that the peptide is not located in the region of the antibody responsible for antigen binding. And importantly, cationic peptides, unlike free and neutral charge peptide-conjugated antibodies, bind to net negative charge ECM. As a result, even though there are 1-2 peptides per antibody, this is sufficient for functional binding to the tumor ECM, which is critical towards studying tumor retention and antitumor efficacy of our cationic peptide-anchored Abs. We subsequently focused on work using cationic peptide as anchors to ICBs to investigate toxicity and efficacy.

### Cationic peptide conjugation reduced systemic exposure of ICBs and showed no treatment-related toxicity

We wanted to confirm the safety of our cationic peptide conjugation to Abs before we investigated their antitumor efficacy. B16F10 murine melanoma cells were inoculated in immunocompetent C57BL/6 mice on day 0. On day 4 after tumor inoculation, mice were randomized into three groups and dosed with one of the following regimens: Abs only (∝-CTLA4 Ab+∝-PD-L1 Ab), Abs-cationic pep (∝-CTLA4–cationic pep+∝-PD-L1–cationic pep), and PBS control (**Fig. 3a**). The peritumorally administered dose was 100 μg (50 ug per Ab), which was consistent with the dose to be used for studying antitumor efficacy. Serum was collected on day 5 and 7 (**Fig. 3a**). The serum concentration of ∝-CTLA4 and ∝-PD-L1 was quantified to measure systemic exposure after peritumoral injection. From serum samples collected on day 5, a significantly lower concentration of ∝-CTLA4 was observed in the Abs-cationic pep group than Abs only group (i.e., free antibody) with a mean of 37.09 and 88.61 μg/mL, respectively (**Fig. 3b**; p<0.0001). On day 7, the concentration of ∝-CTLA4 in both Abs-cationic pep group and Abs only group was lower than day 5 with a mean of 23.60 and 64.75 μg/mL, p<0.0001. The trend was the same for the concentration of ∝-PD-L1 (**Fig. 3c**). The mean rank of ∝-PD-L1 concentration in serum was 28.61 μg/mL in the cationic pep group, compared to 43.56 μg/mL in Abs only group on day 5. On day 7, ∝-PD-L1 concentration in the cationic pep group decreased to 24.38 μg/mL while ∝-PD-L1 concentration in Abs only group dropped to 39.78 μg/mL. The lower serum concentration of both cationic peptide conjugated Abs compared to unmodified Abs suggests that the modification of cationic peptide may help reduce systemic exposure of ICBs.

**Fig. 3.**
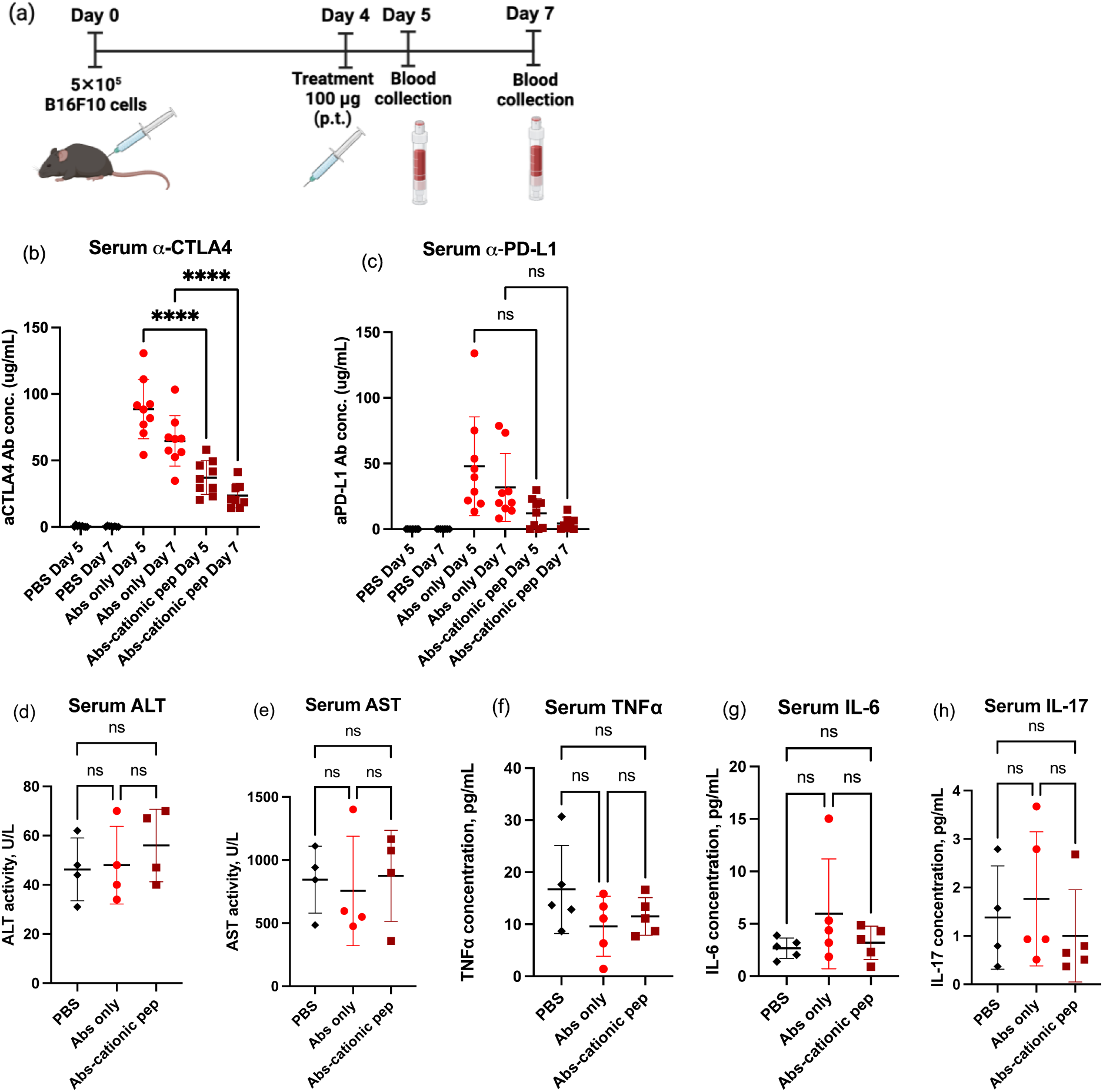
Cationic peptide-anchored ICBs demonstrated decreased exposure in circulation, no treatment-related liver toxicity and no increased release of cytokines after peritumoral injection compared to free ICBs. (**a**) Mice were inoculated with 5 × 10^5^ B16F10 cells on day 0. PBS, Abs only (∝-CTLA4 and ∝-PD-L1 Abs), Abs-cationic pep (∝-CTLA4-cationic pep and ∝-PD-L1-cationic pep) were administered peritumorally on day 4 at 100 µg of Abs (50 µg per each Ab). Blood samples were collected on Day 5 and Day 7. The graphic was generated using BioRender. (**b** and **c**) The concentration of ∝-CTLA4 and ∝-PD-L1 in blood serum was determined by ELISA. n=8 for PBS, Abs-cationic pep; n=9 for Abs only; data pooled from two experimental repeats. (**d** and **e**) Serum ALT and AST activity were measured after blood collection on day 7. n=4; two experimental repeats. (**f** to **h**) Concentrations of TNFα, IL-6, and IL-17 were measured after blood collection on day 7. n=5. All data were shown in mean with SD. Statistical analysis was done using ANOVA with Tukey’s test. Kruskal-Wallis test, followed by Dunn’s multiple comparisons, was used in (b) (c) due to nonparametric data. *P < 0.05, **P < 0.01. ns, not significant.

Next, we studied if ICBs, upon conjugation with the cationic peptide, are well tolerated and measured serum chemistry for activity of liver enzymes alanine aminotransferase (ALT) and aspartate transaminase (AST), along with other analytes, as indicators for hepatotoxicity. Serum samples were collected on day 7 and ALT and AST activities were measured. No statistical significance was observed in either ALT or AST activity amongst the PBS, the cationic pep, and Abs only groups (**Figs. 3d and 3e**). Alkaline phosphatase activity, albumin, and total protein were also measured from the collected serum samples (**Supplementary Information Fig. S1**). No statistical significance was shown amongst the three treatment groups. Collectively, these results indicate that no treatment-related liver damage was observed after a single administration of the cationic peptide conjugated ICBs.

To further investigate systemic toxicity of our cationic peptide conjugated Abs, we measured their ability to induce cytokine and chemokine expression. A panel of inflammatory cytokines and chemokines was assessed with Luminex multiplex assays. Mice serum samples were collected from B16F10 tumor-bearing mice on day 7 after a single peritumoral injection of cationic peptide conjugated Abs, Abs only, or PBS control on day 4. No statistical significance was shown in the concentration of pro-inflammatory cytokines tumor necrosis factor (TNF) α, IL-6, and IL-17 amongst the PBS, the cationic pep, and Abs only groups (**Figs. 3f – 3h**). Besides these three pro-inflammatory cytokines, no statistical differences were found amongst the three groups in any cytokines or chemokines measured from mice serum samples (**Fig. S1**), which suggests that there is an absence of systemic toxicity induced by the peritumoral treatment of Abs. Together, these results demonstrated that cationic peptide conjugation to ICBs reduced systemic exposure compared to unmodified Abs and no treatment-related adverse events was associated with the modified Abs.

### Cationic peptide conjugated ICBs recruited a higher number of activated tumor infiltrating CD8^+^ T cells

We then examined the therapeutic action of peptide-conjugated Abs by characterizing the T-cell responses in melanoma-bearing mice. We inoculated 5× 10^5^ cells in the left flank of C57BL/6 immunocompetent mice on day 0. After which, we administered either PBS or 100 µg of unmodified or peptide-conjugated ICBs (50 µg per Ab) on days 4 and 7 (**Fig. 4a**). After 10 days post tumor inoculation, we isolated and measured leukocytes from tumors and tumor-draining lymph nodes (tdLNs) by flow cytometry (**Fig. 4a**). Cationic peptide conjugated ICBs significantly increased the percentage of CD8^+^CD3^+^ T cells to 22.0±2.5% within the tumor environment compared to 12.3±4.6% for Abs only (**Fig. 4b**). We did not observe any difference in effector population (CD62L^-^CD44^+^) and PD1^+^CD8^+^ T cells until 10 days post tumor inoculation between unmodified and peptide conjugated Abs (**Figs. 4c and 4d**). Similarly, we did not observe any difference between the percentage of CD4^+^CD3^+^ T cells (**Fig. 4e**) and the percentage of effector CD4^+^ T cells (**Fig. 4f**) between PBS, Abs only, and peptide-conjugated Abs. However, we measured a significant reduction in the percentage of CD25^+^Foxp3^+^ T_regs_ in the CD4^+^ population in peptide conjugated ICBs (**Fig. 4g**). There was a 3.9-fold and 1.7-fold increase in the overall CD8^+^ to CD4^+^ T cell population in tumors from the group dosed with cationic peptide conjugated Abs (Abs-cationic peptide) compared to PBS and Abs only, respectively (**Fig. 4h**). Similarly, there was an 8-fold and 1.7-fold increase in CD8^+^ T-cell-to-T_regs_ ratios, which is considered to be predictive of therapeutic efficacy in the B16 melanoma,^40^ observed in the Abs-cationic peptide compared to PBS and Abs only, respectively (**Fig. 4i**). While not statistically significant, we found an average of 2-fold increase in effector to regulatory CD4^+^ population in tumor in cationic peptide conjugated to ICBs compared to PBS (**Fig. 4j**). While we did not notice any difference in effector CD8^+^ T cells, PD1^+^CD8^+^ T cells, and effector CD4^+^ T cells, a significant depletion in the percentage of T_regs_ in the tumor draining lymph nodes (tdLNs) was observed upon cationic peptide conjugation to ICBs (**Fig. 4, k to n**). Similar to the tumor, a significant increase in effector to regulatory CD4^+^ T cell was also found in tdLNs from the group dosed with cationic peptide conjugated Abs (**Fig. 4o**).

**Fig. 4.**
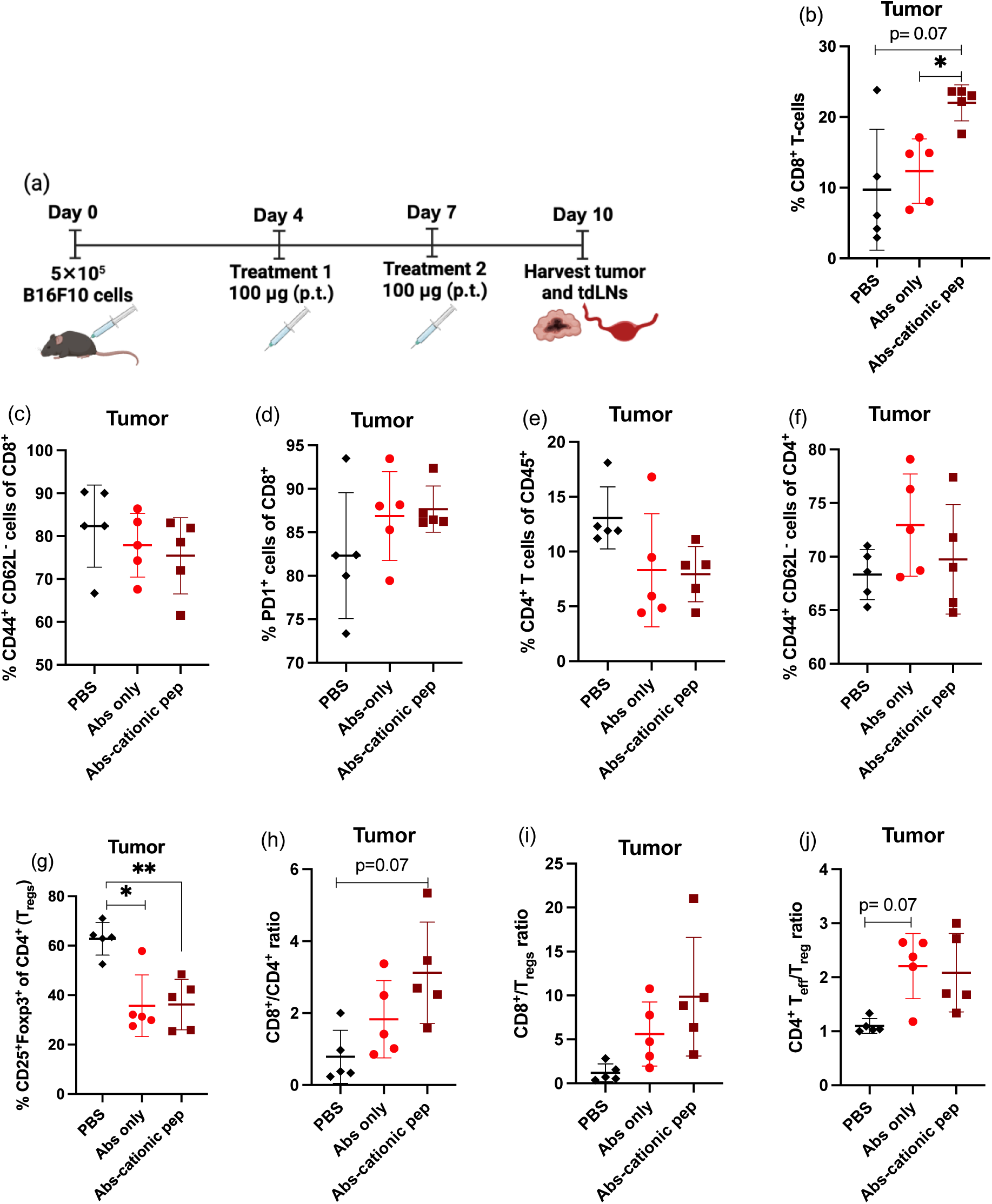
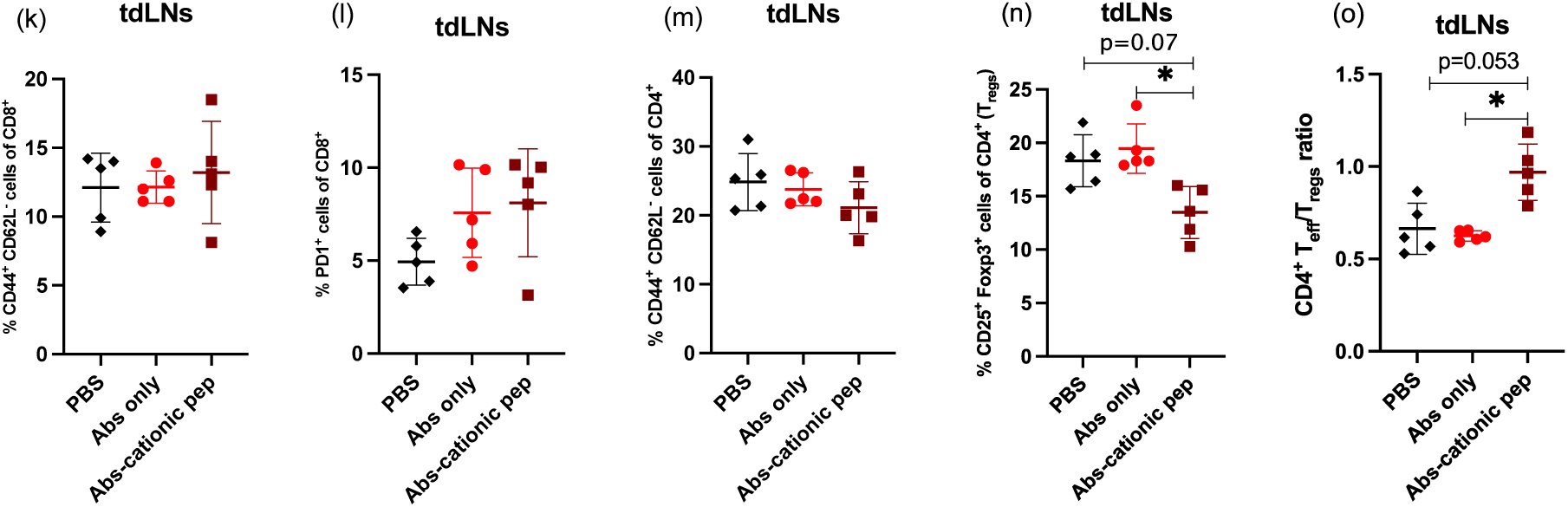
Treatment with cationic peptide-conjugated ICBs increased tumor-infiltrating CD8^+^ T cells and depleted regulatory T cells. (**a**) B16F10 cells were inoculated on day 0. PBS, Abs only (∝-CTLA4 and ∝-PDL1 Abs), or Abs-cationic pep (∝-CTLA4-cationic pep and ∝-PDL1-cationic pep) were administered on days 4 and 7 peritumorally at 100 µg of Abs (50 µg per each Ab). On day 10, tumor and tdLNs were harvested and analyzed for immune phenotyping. The graphic was generated using BioRender. **(b)** frequency of CD8^+^CD3^+^ tumor-infiltrating T cells, **(c)** frequency of CD62L^-^CD44^+^ effector CD8^+^ T cells, **(d)** frequency PD-1^+^ cells among tumor-infiltrating CD8^+^ T cells, **(e)** frequency of tumor-infiltrating CD4^+^CD3^+^ T cells, **(f)** frequency of CD62L^-^CD44^+^ effector CD4^+^ T cells, **(g)** CD25^+^Foxp3^+^ (T_regs_) among CD4^+^ T cells, **(h)** CD8^+^ to CD4^+^ T cell ratio, **(i)** CD8^+^ to T_regs_ ratio, **(j)** T_eff_ to T_regs_ ratio in CD4^+^ T cell, **(k)** frequency of CD62L^-^CD44^+^ effector CD8^+^ T cells in tdLNs, **(l)** frequency of PD-1^+^ cells among CD8^+^ T cells in tdLNs, **(m)** frequency of CD62L^-^CD44^+^ effector CD4^+^ T cells in tdLNs, **(n)** CD25^+^Foxp3^+^ T_regs_ among CD4^+^ T cells in tdLNs, **(o)** T_eff_ to T_regs_ ratio in CD4^+^ T cells in tdLNs. The data was presented as mean±SD (n=5). Statistical analysis was done using ANOVA with Dunnett’s T3 multiple comparisons. *p < 0.05, **p < 0.01.

### Cationic peptide conjugated ICBs suppressed the tumor growth and prolonged survival of melanoma bearing mice compared to unmodified ICBs

Finally, we investigated the antitumor activity of ICBs through combination therapy upon cationic peptide conjugation. B16F10 murine melanoma cells were inoculated in immunocompetent C57BL/6 mice on day 0. Tumored mice were randomized into three treatment groups and dosed with either PBS or 100 µg of unmodified or peptide-conjugated ICBs (50 µg per Ab) on days 4 and 7 after tumor inoculation (**Fig. 5a**). The tumor volume was estimated with time starting day 4. Administration of either unmodified or peptide conjugated Abs significantly delayed tumor growth (**Fig. 5b**). At day 10 of the study, the average tumor volume reduced significantly in the Abs only group (59±48 mm^3^) compared to the PBS group (128±146 mm^3^; p < 0.001). Similarly, the average tumor volume in the Abs-cationic pep was 48±42 mm^3^, which was significantly (p<0.0001) smaller than the PBS group. Although there were no significant differences in tumor volume in Abs-peptide conjugated groups compared to the Abs only group, the average tumor volume was reduced by ∼1.2-fold upon peptide conjugation. We observed an increase in tumor volume for PBS and Abs only groups from day 9 to day 10; interestingly, the tumor volume for the Abs-cationic pep group remained static and did not increase (48±38 mm^3^ on day 9 to 48±42 mm^3^ on day 10). We had to discontinue the tumor regression study when the first mice from Abs only group died and thus only obtained data until day 10 (and not for additional time).

**Fig. 5.**
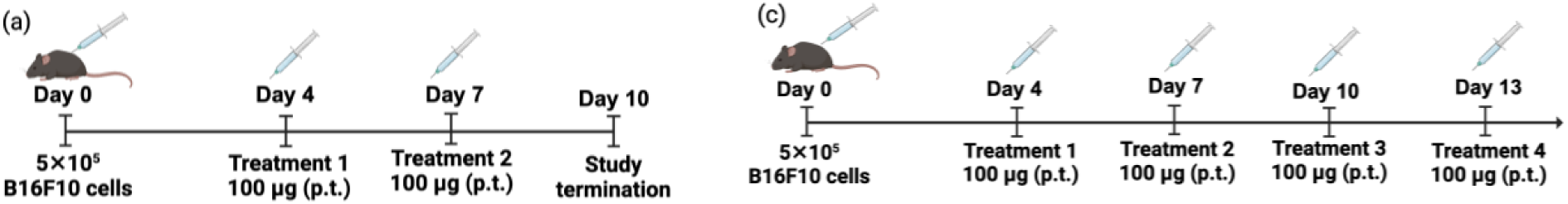
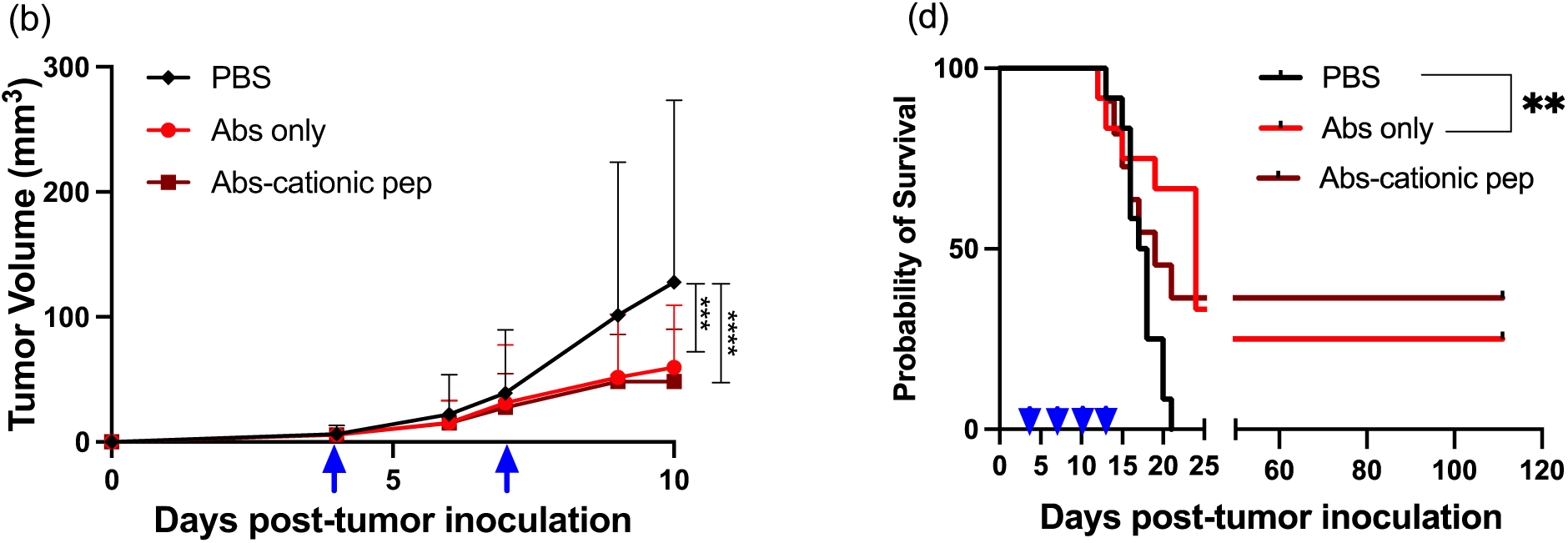
Combination therapy using peptide conjugated Abs (∝-CTLA4 Ab+∝-PD-L1 Ab) reduced B16F10 melanoma growth rate and prolonged survival of tumor-bearing mice. (**a**) B16F10 cells were inoculated intradermally on day 0. PBS, Abs only (∝-CTLA4 Ab+∝-PD-L1 Ab), or Abs-cationic pep (∝-CTLA4–cationic pep+∝-PD-L1–cationic pep) was administered peritumorally on day 4 and 7. Abs dose was maintained at 100 µg (50 µg of each ∝-CTLA4 Ab and +∝-PD-L1 Ab) per dose. (**b**) Tumor volumes are presented as means ± SD; n= 17 for PBS and Abs only; n=16 for Abs-cationic pep; data pooled from two experimental repeats; statistical analyses were done using ANOVA with Turkey’s multiple comparisons test. Graphs depict tumor volume until the first mouse died. (**c**) B16F10 cells were inoculated intradermally on day 0. PBS, Abs only (∝-CTLA4 Ab+∝-PD-L1 Ab), or Abs-cationic pep (∝-CTLA4–cationic pep+∝-PD-L1– cationic pep) was administered peritumorally on day 4, 7, 10 and 13 (**d**) Probability of survival is plotted with days post tumor inoculation; n= 12 for PBS and Abs only; n=11 for Abs-cationic pep. Statistical analyses were done using the log-rank (Mantel-Cox) test. *p < 0.05, **p < 0.01, ***p < 0.001, ***p < 0.0001. The graphic was generated using BioRender.

Next, we validated whether the administration of cationic peptide conjugate could prolong the percent survival in the melanoma model. Here, the tumored mice were randomized into three treatment groups and dosed with either PBS or 100 µg of unmodified or peptide-conjugated ICBs (50 µg per Ab) on days 4, 7, 10 and 13 after tumor inoculation (**Fig. 5c**). The body weight of the mice in each group remained within an acceptable range for the duration of the study, which indicates that the treatment regimen of Abs and Abs-peptide conjugates were well tolerated by the mice (**Fig. S2**). Administration of 100 µg of either Abs alone or Abs conjugated to the peptides prolonged the percent survival compared to the untreated PBS group (**Fig. 5d)**. The median survival was increased from 17.5 days for the PBS group to 24 and 19 days for Abs only and Abs-cationic pep treatment groups, respectively. Similarly, when 100% of mice from the PBS group had died within 21 days from the tumor inoculation, the survival population of Abs only and Abs-cationic pep groups after 110 days of tumor inoculation were 25% and 37%, respectively. The slower increase in tumor volume for the Abs-cationic pep group and a relatively higher overall percent survival suggest cationic peptide has the potential to delay tumor growth compared to free, unconjugated Abs. The apparent difference between Abs only and Abs-cationic peptide groups in the survival study (but not in the tumor regression) can be explained by the heterogeneous treatment response observed (**Fig. S3**).

## DISCUSSION

Local administration of ICBs has been considered an attractive alternative to overcome the challenges due to systemic administration.^28^ Local immunotherapy limits the systemic exposure and allows for lower dose administration, yet still generates systemic antitumor response of the immunotherapies. Here, we have leveraged electrostatic interactions with the tumor ECM to enhance the retention of ICBs in the microenvironment of murine melanoma. Contrary to the traditional approach with neutral charge surfaces, we had previously identified a cationic peptide that enhanced the uptake, penetration, and retention of a biological nanoparticle in tumor tissue compared to a neutral peptide.^30^ We had discovered the neutral peptide using phage display, which had minimal electrostatic interactions with the charged components of the tumor ECM.^41^ Whereas, the cationic peptide interacted electrostatically with the negatively charged hyaluronic acid of the tumor ECM.^30^ Based on these insights, we conjugated the cationic peptide to ∝-CTLA4 and ∝-PD-L1 ICB antibodies to evaluate whether the electrostatic interaction-mediated intratumoral retention of these Abs can generate positive, antitumor immunotherapeutic responses.

We conjugated the cationic and the neutral peptide (as a control) to the ICBs using copper-free click chemistry. We estimated an average of two of each peptide were conjugated to the ∝-CTLA4 Ab and one of each peptide were conjugated to the ∝-PD-L1 Ab. Regina et al. followed similar copper-free click chemistry to conjugate four Angiopep-2 peptides to ∝-HER2 monoclonal antibodies, as calculated using matrix-assisted laser desorption/ionization-time-of-flight (MALDI-TOF) mass spectrometry.^42^ The differences in the number of conjugated peptides might be due to the number of free amine groups available per different types of monoclonal antibodies. For example, Ishihara et al., through sulfosuccinimidyl-4-(N-maleimidomethyl) cyclohexane-1-carboxylate (sulfo-SMCC) mediated chemistry, covalently linked placenta growth factor-2 (PIGF-2123-144) collagen-binding peptide to ICBs and reported an average of 6.3 and 6 PIGF-2123-144 peptides were bound to monoclonal ∝-CTLA4 and ∝-PD-L1 Abs.^29^ These findings suggest that changing and optimizing conjugation chemistry can increase the valency of peptides per antibody.

Next, we demonstrated that the peptide conjugation did not hinder the antigen recognition and binding ability of the ICBs. Yet only the cationic peptide conjugated Abs bound to the net negative charge ECM, unlike free and neutral charge peptide-conjugated Abs. The neutral peptide conjugated ∝-CTLA4 Ab did not bind to the tumor ECM, and the neutral peptide conjugated ∝-PD-L1 Ab exhibited weak binding to the tumor ECM. In contrast, the cationic peptide conjugated Abs bound to tumor ECM with stronger binding affinities than neutral peptide conjugated Abs despite having similar conjugation efficiency (∼two peptides per ∝-CTLA4 Ab and ∼one peptide per ∝-PD-L1 Ab). The binding behavior of ICBs conjugated with cationic and neutral charge peptides correlates with our previous studies studying retention of phage displaying cationic or neutral charge peptides in tumor ECM and tumor tissue. We did observe the neutral peptide having minimal electrostatic interactions with the charged components of the tumor ECM.^41^ Contrary, we observed that phage displaying cationic peptide interacted electrostatically with the negatively charged hyaluronic acid of the tumor ECM.^30^ Therefore, similar to our prior work, the enhanced ECM affinity of cationic peptide conjugated Abs is due to the electrostatic binding of the cationic peptide with the negatively charged hyaluronic acid present in the tumor ECM. From Ishihara et al, PIGF-2_123-144_, upon conjugation to ICBs, had stronger ECM binding affinity (k_d_ of 1.3 nM for PIGF-2_123-144_ conjugated to ∝-CTLA4 Ab, and 5.1 nM for PIGF-2_123-144_ conjugated to ∝-PD-L1 Ab) compared to our cationic peptide conjugated ICBs.^29^ PIGF-2_123-144_ peptide was multivalently conjugated on Abs (∼6 peptides per Ab), which may have contributed to the binding. Regardless, our cationic peptides achieve binding without relying on the need for multivalent interactions (∼1-2 peptide per Ab).

We further investigated that a single peritumoral administration of 100 μg of cationic peptide conjugated ICBs may help reduce the systemic exposure of ICBs, without indicating any treatment-related adverse events. Our results using charge-based peptide conjugates align with other work where increasing the binding of locally administered immunotherapies to tumor ECM reduced systemic exposure. Work has shown that ∝ −CTLA4 and ∝ −PD-L1 ICB Abs conjugated with peptide PIGF-2_123-144_, which binds multiple ECM proteins, exhibited lower concentrations in blood plasma compared to unmodified equivalents after peritumoral injection into B16F10 tumors.^29^ Interestingly, similar to our findings, the concentration of ∝ −PD-L1 Abs with or without PIGF-2_123-144_ modification in blood plasma was much lower than the concentration of corresponding ∝ −CTLA4 Abs. In addition to conjugation to ICBs, PIGF-2_123-144_ was conjugated to ∝-CD40 antibody agonist and observed similar behavior after peritumoral injection into B16F10 tumors. The concentration of ∝ −CD40 in blood serum was lower in PIGF-2_123-144_ modified ∝ −CD40 arm than ∝ −CD40 arm, which corroborated that PIGF-2_123-144_ helped reduce systemic exposure of these immunotherapies.^43^ In addition to PIGF-2_123-144_, collagen-binding protein lumican was used to increase binding of a mouse serum albumin fusion protein to tumor ECM after intratumoral injection into B16F10-TRP2KO tumors, which prolonged retention of the albumin fusion in tumors while decreasing the concentration in serum.^44^ With these collective findings, they support that conjugation with our cationic peptide may help reduce systemic exposure by increasing binding and retention to the net negatively charged tumor ECM, as shown earlier (**Fig. 2**).

Next, we measured cytokine and chemokine expression after administration of our cationic peptide conjugated and non-conjugated ICBs. Proinflammatory cytokines released into circulation during ICB therapy may serve as potential biomarkers to identify individuals at risk of developing immune-related adverse events to ICB.^45^ In a study conducted on advanced melanoma patients treated with either ∝-PD-1 monotherapy (pembrolizumab or nivolumab) or ∝-PD-1 in combination with ∝-CTLA-4 (ipilimumab), it was found that a panel of 11 cytokines, including G-CSF (Granulocyte-Colony Stimulating Factor), IL-1α, IL12p70, and IL13, were upregulated in patients with severe immune-related adverse events preceding and during treatment relative to patients without severe immune-related adverse events.^46^ In a study of 35 melanoma patients who received ∝ −CTLA4 Ab ipilimumab, there was higher blood circulating IL-17 level compared to baseline, which was significantly associated with grade 3 diarrhea/colitis.^47^ In another study of ∝ −CTLA4 Ab ipilimumab treated metastatic melanoma patients who displayed ICB-induced colitis presented elevated serum IL-17 concentrations after treatment compared to patients who did not have immune-related adverse events.^48^ In addition to IL-17, induction of IL-6 has been shown to be associated with immune-related adverse events development including psoriasiform dermatitis and colitis.^49, 50^ Of note, we evaluated the production of proinflammatory cytokines and chemokines after treatment with cationic peptide conjugated Abs and found no significant difference relative to either the PBS group or unmodified Abs group, which suggests no treatment-related adverse events would be expected from the release of cytokines and chemokines due to the treatment or the modification of Abs with our cationic peptide. Along with these findings, there are no changes in body weight of the tumored mice when dosed with cationic peptide conjugated ICBs (**Fig. S2**), these findings collectively indicate that conjugation with our cationic peptide anchor is well tolerated and nontoxic upon local administration.

Blockade of PD-L1/PD-1 and CTLA4 negative costimulatory pathways allows tumor-specific T cells to expand and carry out effector functions that otherwise would have been inactivated.^5^ Therefore, the combinatorial blockade shifts the tumor microenvironment from suppressive to inflammatory. Curran et al. reported combination blockade of the CTLA4 and PD-1 pathways strongly increased infiltration of CD8^+^ T cells in melanoma environment.^5^ When PD-L1 blockade alone modestly increased T-cell infiltration, CTLA4 blockade alone failed to show any effect. However, they found an increase in CD8^+^ T-cell-to-T_regs_ ratios within tumors upon CTLA4, PD-1, or PD-L1 blockade. They also reported significant improvement in CD4^+^ T_eff_/T_regs_ ratios within tumors upon only CTLA4 blockade and further improvement using combination therapy of ∝-CTLA4, ∝-PD-1, and ∝-PD-L1 compared to monotherapies in melanoma.^5^ Ishihara et al. coupled an ECM binding peptide (PIGF-2_123-144_) with ICBs (∝-CTLA4 and ∝-PD-L1) to improve the retention of ICBs upon local administration.^29^ They treated B16F10 melanoma with 200 µg (100 µg per Ab) of unmodified or peptide-conjugated Abs on 4 and 7 days post tumor inoculation and extracted the leukocytes from tumor and tdLNs on day 8. They reported an increase in the percentage of CD8^+^ T cells in the tumor environment upon increasing the retention of ICBs in the tumor by peptide conjugation. They reported an increase in effector population and PD-1^+^CD8^+^ T cells compared to PBS treatment. While we did not determine a statistical difference in the effector and PD-1^+^ population in CD8^+^ T cells, the percentage of PD-1^+^CD8^+^ T cells increased from 82.3±7.2% in PBS to 87.7±2.6% in the Abs-cationic pep group in the tumor environment. Unlike Ishihara et al., although we did not see any difference in the percentage of effector CD4^+^ T cells among study groups, we observed a 2.0-fold increase in effector to regulatory CD4^+^ T cell ratios in tumor environment upon cationic peptide conjugation to ICB Abs compared to PBS. The anomaly observed between these two studies might be due to the two-fold increased dosing of Abs (200 µg per injection) used by Ishihara et al. compared to our dose (100 µg per injection).

While Ishihara et al. reported a similar percentage of T_regs_ among all groups (including PBS),^29^ we found a significant depletion of T_regs_ in the tumor environment upon administration of either unmodified or peptide-conjugated ICBs compared to PBS (**Fig. 4g**). As we used the equivalent ∝-CTLA4 clone (9H10) and ∝-PD-L1 clone (10F.9G2) in our study as used by Ishihara et al., the inconsistencies in this observation might be because of the differences in the time scale of immune cell phenotyping. While we measured the immune cell population on day 10, Ishihara et al. performed their study after 8 days post-tumor inoculation. ∝-CTLA4 clone (9H10) has been investigated to deplete T_regs_ in tumor microenvironments previously.^24, 51, 52^ Duraiswamy et al. found that PD-1 or PD-L1 blockade alone reduced T_regs_ moderately in the tumor, while CTLA4 blockade alone (using clone 9D9 Ab) was not able to affect T_regs_ numbers.^15^ However, combined blockade of CTLA4 and PD-1/PD-L1 pathway further reduced T_regs,_ which resulted in increased CD8^+^/T_regs_ and CD4^+^/T_regs_ ratios.^15^ They found that T_regs_ were primarily comprised in the CTLA4^+^CD4^+^ population, and among the CTLA4^+^ cells, most were Foxp3^+^CD25^+^ cells, representing activated effector T_regs_. PD-L1 plays a pivotal role in regulating induced T_regs_ cell development.^40^ Therefore, blocking CTLA4 and PD-L1 simultaneously by cationic peptide-conjugated ICBs leads to a reduction in T_regs_ population (**Fig. 4g**) and an increase in CD8^+^/T_regs_ (**Fig. 4i**) and CD4^+^/T_regs_ ratios (**Fig. 4j**). Similarly, we also found a significant reduction in T_regs_ (**Fig. 4n**) and a significant increase in CD4^+^/T_regs_ ratios in tdLNs upon cationic peptide conjugation (**Fig. 4o**). The cationic peptide increased the binding of the ICBs to the negatively charged tumor ECM electrostatically, as shown in Fig. 2. The increased binding likely improved the retention of the Abs in the tumor environment, which resulted in increased infiltration of activated CD8^+^ T cells and depletion in immune suppressive T_regs_ in tumor and tdLNs. The increased population of cytotoxic CD8^+^ T cells potentially produced higher levels of effector cytokines, which we speculate would improve the therapeutic efficacy of ICBs.

Monotherapy of ∝-CTLA4 and ∝-PD-L1 Abs has been FDA-approved to treat melanoma. ∝-CTLA4 Ab was administered to patients with metastatic melanoma at a dose of 3 mg per kilogram of body weight through intravenous infusion along with a gp100 peptide vaccine.^8^ Similarly, ∝-PD-L1 Ab was administered at a dose of 0.3 to 10 mg per kilogram of body weight to patients with selected advanced cancers, including melanoma.^53^ Combinatorial blockade of CTLA4 and PD-L1 resulted in additive or synergistic therapeutic effects against melanoma tumor models. In a subcutaneous B16.SIY melanoma tumor model, Spranger et al. observed significant tumor regression with the combination of ∝-CTLA4 and ∝-PD-L1 compared to monotherapy with either Ab.^2^ ∝-CTLA4 Ab was administered at 100 µg per mouse i.p. at three single time points (day 4, 7, and 10), and ∝-PD-L1 Ab was administered at 100 µg per mouse i.p. every other day starting day 4 and ending on day 16 post tumor inoculation. Compared to single regimens, the combinatorial treatment of ∝-CTLA4 and ∝-PD-L1 improved tumor control, with 15 complete responders observed out of 27 treated mice.^2^ Although combination therapy of ∝-CTLA4 and ∝-PD-L1 Abs has promising therapeutic efficacy in the clinic for various types of cancer,^4, 6, 7^ there are systemic toxicity and immune-related adverse events from treatment that greatly limit efficacy.^53, 54^

To overcome the challenges associated with systemic administration, Francis et al. administered 150 µg each of a ∝-CTLA4 and ∝-PD-1 Ab i.p. or i.t. (300 µg total, three times the dose used in our study) on days 5, 7, and 9 after tumor implantation.^24^ They observed a significant delay in tumor growth and improved overall survival in groups locally dosed via i.t administration compared to groups dosed with untreated PBS control and i.p. administration.^24^ However, Ishihara et al. reported that the conventional combination therapy of ∝-CTLA4 and ∝-PD-L1 administered i.p. or peritumorally (p.t.) displayed insufficient therapeutic efficacy compared to the PBS control; conjugating an ECM binding peptide enhanced the retention of the Abs in the tissue which substantially delayed growth in melanoma models. They varied the dose of the Abs starting from 25 µg to 300 µg of each Ab, which were administered every three days for three doses starting on day 4 after the tumor implantation. Upon 200 µg administration (twice that used in our study) of the peptide conjugated ICBs, the tumor size measured after 9 days of tumor inoculation is similar to what we found in our study, i.e., the average volume of ∼30-50 mm^3^. However, while we observed similar tumor regression in the unmodified Abs group (with an average tumor volume of 51 mm^3^), Ishihara et al. reported a larger average tumor size for their unmodified group on day 9 (∼150 mm^3^).

Here, we demonstrate that the addition of cationic peptides as anchors can enhance the therapeutic efficacy of immune checkpoint blockade antibodies, yet it is feasible to improve upon this charge-based anchoring approach. In our work, we conjugated an average of 1 – 2 peptides per antibody using click chemistry. Even at the low valency of peptides, we achieved strong binding to the ECM without compromising the antigen binding of the antibodies. Also, there was enhanced infiltration of activated CD8^+^ T cells in the tumor environment and suppression of T_regs_ in both tumor and tdLNs. We posit that improving the number of peptides per antibody by changing the conjugation chemistry might strengthen the avidity of electrostatic interactions between the peptide-conjugated antibodies with the tumor environment for improved retention and antitumor efficacy. However, it is feasible to achieve efficacy with low valency, as work with lumican fusion proteins, whereby one collagen-binding peptide was fused to therapeutic cytokines through genetic engineering, improved delay in tumor growth and antitumor efficacy in different models of cancer.^44^

It may also be necessary to optimize the charge of the peptide to have more favorable interactions with the melanoma tumor microenvironment for improved uptake, penetration, and retention of anchored antibodies. In our previous work, we identified and confirmed the enhanced penetration and retention of nanoparticles displaying cationic peptides within a pancreatic tumor environment.^30^ The human pancreatic BxPC3 environment (0.27±0.04 mg/g) has a more negative fixed charge density (i.e., net charge of environment) compared to murine melanoma B16F10 environment (0.006±0.001).^36^ This is due to the greater negative charge tumor ECM of pancreas tumors compared to melanoma tumors due to the presence and abundance of hyaluronic acid. As a result, cationic peptides of the same charge will have different strengths of electrostatic interactions with different tumor microenvironments. The interaction between the cationic peptide and a tumor microenvironment is context-dependent on the charge of the microenvironment. The charge of the cationic peptide can be optimized based on the fixed charge density of the tumor environment to favorably harness the electrostatic interaction for enhanced retention and penetration to ultimately improve therapeutic outcomes of cancer therapies.

## MATERIALS & METHODS

Hamster anti-mouse CTLA4 clone 9H10 antibody (∝-CTLA4, #BP0131) and rat anti-mouse PD-L1 antibody clone 10F.9G2 (∝-PD-L1, #BP0101) along with their respective dilution buffers InVivoPure pH 7.0 dilution buffer (#IP0070) and InVivoPure pH 6.5 dilution buffer (#IP0065), and their corresponding isotype controls, InVivoPlus polyclonal Syrian hamster IgG (#BP0087) and InVivoPlus rat IgG2b isotype control (Clone LTF-2, #BP0090) were purchased from BioXCell. Click-easy mono fluoro-substituted cyclooctyne (MFCO) (#LK4300) was purchased from Berry & Associates, Inc. PD-10 desalting column containing Sephadex G-25 medium (#GE17-0851-01) was from GE healthcare, and Amicon Ultra-4 centrifugal filter devices, 50K MWCO (#UFC805024) was purchased from EMD Millipore Sigma. Azide-modified neutral peptide (Lys(Azide)-CKPGDGGPC, M.W. 985.12 gm/mol) and Azide-modified cationic peptide (Lys(Azide)-CRRRRKSAC, M.W. 1287.55 gm/mol) were synthesized following Fmoc chemistry by LifeTein, LLC. Float-A-Lyzer G2 dialysis devices, 50K MWCO (#G235034), were purchased from Spectrum Laboratories, Inc. For reduced SDS-PAGE, 10X Tris/Glycine/SDS buffer (#1610732), 10% Mini-PROTEAN® TGX™ Precast Protein Gels (#456-1036), 4× Laemmli sample buffer (#1610747), and Precision Plus Protein™ Dual Xtra Prestained Protein Standards (#1610377) were purchased from Bio-Rad. 10× bolt sample reducing agent, 500 mM dithiothreitol (DTT) (#B0009), SimplyBlue^TM^ SafeStain (#LC6060), Nunc Maxisorp^TM^ flat-bottom ELISA 96-well plate (#44240421), and 1-Step™ Ultra TMB-ELISA Substrate Solution (#34028) were procured from Thermo Fisher Scientific. We purchased recombinant mouse CTLA4 Protein (rmCTLA4, #50503-M08H) and PD-L1 Protein (rmPD-L1, #50010-M08H) from SinoBiological. Horseradish Peroxidase (HRP)-labeled secondary Abs; Rabbit Anti-Rat IgG H&L (HRP) (#ab6734) and Rabbit Anti-Syrian Hamster IgG H&L (HRP) (#ab6783) were purchased from Abcam. Hyaluronic acid (HA) sodium salt extracted from rooster comb (#9067-32-7) was obtained from Sigma, and rat tail collagen Type I (#354249) was obtained from Corning. SST-MINI serum separator tubes (#450571VET) were provided by IDEXX BioAnalytics. Foxp3/Transcription factor staining buffer set (#005523) and cell stimulation cocktail (#00497093) were purchased from eBioscience. eBioscience™ fixation/permeabilization concentrate (#00512343), eBioscience™ fixation/permeabilization diluent (#00522356), eBioscience™ permeabilization buffer (#00833356), Iscove’s Modified Dulbecco’s medium (#12440046), Falcon™ Cell Strainers (#087712) were purchased from Thermo Fisher Scientific. Collagenase D (#11088866001), DNAse I (#DN25-100MG), sodium azide (#S2002-25G) were from Milipore Sigma. Brefeldin A Solution (#420601), purified anti-mouse CD3e antibody (#100302), purified anti-mouse CD28 antibody (#102102), fixation buffer (#420801), and intracellular staining permeabilization wash buffer (#421002) were purchased from BioLegend.

### Synthesis of antibody peptide conjugates

We synthesized antibody peptide conjugates through copper-free click chemistry following a two-step synthesis process as previously described with some modifications.^42^ In the first step, the antibodies were modified to incorporate linker moieties to the antibodies, while in the second step, azide-modified peptides were added to form an azide-alkyne bond (**Fig. 1a**). Briefly, in step 1, Ab stocks were diluted to 3 mg/mL in their respective dilution buffers. The Abs (1-3 mL of 3 mg/mL) were structurally modified by incubating with 12 equivalents of MFCO solution dissolved in DMSO (at a stock concentration of 25 mg/mL) for 6 h at RT while shaking at 2 rpm. The modified Abs were purified from other reagents by gel filtration using PD-10 desalting columns, followed by buffer exchange using Amicon filter (50K MWCO) to resuspend the modified Abs in their respective dilution buffers. In step 2, 8 equivalents of azide-modified peptides resuspended in DMSO were incubated with the modified Abs overnight at RT while shaking at 2 rpm. Following, the Ab-peptide conjugates were purified using dialysis against the respective dilution buffers using 50 kDa MWCO float-a-lyzer at 4 °C for 24 h. For the Ab-only group, synthesis steps were carried out in parallel, where equal volumes of DMSO were added instead of MFCO or peptide in step 1 and step 2, respectively.

#### Sodium Dodecyl Sulfate-polyacrylamide gel electrophoresis (SDS-PAGE)

We verified the conjugation of peptides to the ICBs following SDS-PAGE under reduced conditions using the Mini-PROTEAN Tetra system (Bio-Rad). The unmodified and peptide-conjugated ICBs were reduced to corresponding heavy and light chains of the Abs by incubating the formulations with 50 mM DTT for 10 min at 70 °C. Reduced formulations were then loaded to a 10% polyacrylamide gel at 3 µg/lane of Ab, and the gel was run at 100 V for 1 h. Simultaneously, 10 µL of the pre-stained protein standard was also run through the gel to mark the molecular weight. After gel electrophoresis, the gel was stained with Coomassie blue following the SimplyBlue SafeStain protocol. Further, the gels were imaged using Syngene G:Box gel doc system.

### Liquid Chromatography-Mass Spectrometry (LC-MS)

We estimated the number of peptides conjugated per antibody using LC-MS. Unmodified or peptide-conjugated Abs were reduced using DTT to quantify the number of peptides per heavy and light chain of each Ab. 20-30 µL of 1 mg/mL formulations were incubated with 50 mM DTT for 10 min at 70 °C. The molecular weight of the reduced Abs was analyzed by the University of Texas at Austin Proteomics Facility (Center for Biomedical Research Support). The samples were analyzed using an Orbitrap Fusion mass spectrometer using the ion trap detector and deconvoluted with Thermo Protein Deconvolution software.

### Tumor-like ECM model preparation

As established in our previous studies, we prepared a tumor-like ECM model consisting of collagen and hyaluronic acid (HA).^30, 41^ Briefly, HA was suspended in phosphate-buffered saline (PBS) 1× while stirring at 350 rpm at 4 °C to prepare a 5-7 mg/mL stock solution. The solution was stirred continuously for 10 h and left overnight at 4 °C. Collagen stock (8-10 mg/mL) was pH neutralized, and its ionic strength was maintained following the manufacturer’s protocol. Then, the HA solution was spiked into the neutralized collagen while stirring at 125 rpm at 4 °C for 10 min. The final concentration of collagen and HA were maintained at 7.8 mg/mL and 0.84 mg/mL, respectively, to mimic the tumor ECM.

### Quantification of Ab binding affinity to ECM proteins and recombinant proteins using ELISA

First, we diluted the tumor-like ECM, recombinant mouse (rm) rmCTLA-4, and rmPD-L1 in carbonate buffer as the coating buffer (0.1M Na_2_CO_3_-NaHCO_3_, pH 9.0). The tumor ECM concentration was maintained at collagen and HA concentrations of 10 µg/mL and 1.05 µg/mL, respectively. Similarly, the concentration of rmCTLA-4 and rmPD-L1 were diluted to 1 µg/mL. Then, the proteins were coated in Maxisorp ELISA 96-well plates at a volume of 100 µL/well and incubated at 4 ℃ overnight. The following day, the coating buffer was removed, and the wells were washed five times using 300 µL/well of washing buffer (PBS with 0.05% Tween 20). Next, the wells were blocked by incubating with 200 µL/well of blocking buffer (2% BSA w/v in washing buffer) for 2 h at RT. Meanwhile, the unmodified and peptide-conjugated ICBs were diluted serially in the blocking buffer. Next, the blocking buffer was removed from the plate, and 100 µL of samples were added per well to bind with the proteins without washing the wells. The samples were incubated at RT for 2 h. Then the samples were removed, and the wells were washed three times with the washing buffer at a volume of 300 µL/well. Next, the respective HRP-conjugated secondary Abs (5000-fold dilution in 4 times diluted blocking buffer) were incubated for 1 h at RT. The plates were again washed five times with washing buffer after removing the secondary Abs. Next, 100 µL of TMB was added to each well and incubated at RT in the dark for 15-30 m. The reaction was stopped by adding 100 µL of 2M sulfuric acid into each well. The absorbance was read at 450 nm using a plate reader. The binding dissociation constant (𝐾_𝑑_) was estimated by non-linear least square regression analysis in excel, assuming one-site-specific binding. The theoretical 𝐾_𝑑_ was calculated using absorbance at 450 nm, 𝐴_450_ = 𝐵_𝑚𝑎𝑥_ × [𝐸𝐶𝑀_𝑝𝑟𝑜𝑡𝑒𝑖𝑛_⁄(𝐾_𝑑_ + 𝐸𝐶𝑀_𝑝𝑟𝑜𝑡𝑒𝑖𝑛_)].

### Mice and cell lines

C57BL/6 female mice, ages 8-12 weeks, were obtained from either Taconic Biosciences or Jackson Laboratory. Experiments were performed in accordance with the United States Public Health Service Policy on Humane Care and Use of Laboratory Animals. The animal protocols were approved by the University of Texas at Austin Institutional Animal Care and Use Committee (IACUC). B16F10 cells were obtained from the American Type Culture Collection (ATCC) and cultured as per instructions with some required modifications. Briefly, ∼2× 10^6^ cells were seeded in T175 flask with 35 mL Dulbecco’s Modified Eagle’s Medium (DMEM) supplemented with 10% Fetal Bovine Serum (FBS) and 1% Penicillin-Streptomycin (Pen-Strep). After three days, the media was changed, and the following day the cells were passaged. We observed B16F10 cells to be highly sensitive to trypsinization; therefore, for the cell detachment, we incubated the cells with trypsin for 2.5 m max at 37 °C in a 5% CO_2_ incubator. After three more days, the cells were prepared for inoculation.

### In vivo toxicity

A total of 5× 10^5^ cells resuspended in 50 µL of DMEM were injected intradermally on the left flank of each mouse. After 4 days, mice were injected peritumorally (intradermally beside the tumor) with 100 μg (50 μg per Ab) of unmodified or cationic peptide conjugated ICBs. Equivalent volume of PBS was injected peritumorally as control group. Blood samples were collected by tail snip in SST-MINI serum separator tubes 5 days after tumor inoculation. On day 7, mice were humanely sacrificed. Blood samples were collected by cardiac puncture and transferred into serum separator tubes. Serum samples were separated by centrifugation after the blood was clotted. Concentrations of ICBs in serum were measured by ELISA as described above. Serum samples were sent to IDEXX BioAnalytics for liver panel testing. Serum samples were sent to Eve Technologies for measurement of cytokines and chemokines (Mouse Cytokine Array/Chemokine Array 32-Plex panel).

### Tissue harvesting and single-cell suspension preparation

A total of 5× 10^5^ cells resuspended in 50 µL of DMEM were injected intradermally on the left flank of each C57BL/6 mouse. After 4 and 7 days, 100 µg (50 µg per Ab) of unmodified or cationic peptide conjugated ICBs, along with the untreated PBS control, were injected peritumorally. Mice were sacrificed on day 10; tumor and tumor-draining lymph nodes (inguinal lymph nodes, tdLN) were harvested. Tumors were collected in a collection buffer consisting of DMEM and 1× Pen-Strep, whereas tdLNs were collected in 1× PBS. Tumors were first finely minced in a weighing boat and then digested in 2.5 mL of digestion buffer (DMEM supplemented with 2% FBS, 1× Pen-Strep, 2 mg/mL collagenase D, and 40 µg/mL deoxyribonuclease I) for 30 min at 37 °C, while shaking gently at every 10 min interval. Tumor digestion was quenched with DMEM + 20% FBS. Single-cell suspensions of tumors (after incubation) and tdLNs were obtained by gently disrupting the organs to push them through a 70-mm cell strainer. The strainer was washed with the flow-through and spun down at 200 ×g for 5 min at 4 °C. The pellet was resuspended in cold PBS and washed two times in PBS. The cells were spun down for 5 min at 300 ×g, and the pellet was resuspended in a freezing buffer (50% RPMI, 40% FBS, and 10% DMSO) and froze under liquid N_2_ until ready for further analysis.

### Ab staining and immune cell phenotyping

The single-cell suspensions of tumor and tumor-draining lymph nodes (tdLNs) were stained with Abs for different cell surface markers, intracellular Foxp3, and cytokines. Abs against the following molecules were used for staining: CD45 (30-F11, #103108, BioLegend), CD3 (145-2C11, #100330, BioLegend), CD4 (RM4-5, #100511, BioLegend), CD8 (53-6.7, #100738, BioLegend), CD44 (IM7, #740215, BioLegend), CD62L (MEL-14, #612833, BD Biosciences), CD25 (PC61, #102035, BioLegend), Foxp3 (MF14, #560401, BioLegend) and PD-1 (29F.1A12, #135225, BioLegend). eBioscience Fixable Viability Dye eFluor 506 (#65-0866-1, eBioscience) was used for the viability stain. Frozen cells were thawed and resuspended in FACS Azide buffer (0.05% sodium azide in 500 mL PBS, 50 mM EDTA, and 2% FBS). The cells were then stained with fixable viability dye and cell surface marker cocktail (1 µL from each dye) for 45 mins by incubating at 4 °C in the dark. The cells were then washed two times with 150 µL FACs Azide buffer. Next, the intracellular staining was performed using a Foxp3 staining kit according to the manufacturer’s protocol. According to this, 200 µL of Foxp3 fixation/permeabilization working solution was incubated with the cells for 30-60 min at RT in the dark. Then the cells were washed two times with 200 µL of dye conjugated Ab for detecting intracellular antigen Foxp3 was incubated for at least 30 min at RT in the dark. The cells were then washed two times with 200 µL of 1× permeabilization buffer. The stained cells were resuspended in 500 µL of FACs Azide buffer for further immune cell phenotyping using flow cytometry. All flow cytometric analyses were performed using a BD FACSAria Fusion SORP cell Sorter and analyzed using FlowJo software.

### Tumor regression study

A total of 5× 10^5^ cells resuspended in 50 µL of DMEM (without FBS) were injected intradermally on the left flank of each mouse. After 4 and 7 days, 100 µg (50 µg per Ab) of unmodified or peptide conjugated ICBs, along with the untreated PBS control, were injected peritumorally (intradermally beside the tumor). The study groups were PBS, Abs only (∝-CTLA4 and ∝-PDL1 Abs), or Abs-cationic pep (∝-CTLA4-cationic pep and ∝-PDL1-cationic pep). Tumors were measured with a digital caliper starting day 4 (when the tumors were visible), and the volume of the tumors was tracked with the number of days. We estimated the volume based on the length and width of tumors as 𝑉 = (1⁄2) × 𝑙𝑒𝑛𝑔𝑡ℎ × 𝑤𝑖𝑑𝑡ℎ^2^.

### Overall survival study

A total of 5× 10^5^ cells resuspended in 50 µL of DMEM were injected intradermally on the left flank of each mouse. After 4, 7, 10, and 13 days, 100 µg (50 µg per Ab) of unmodified or peptide conjugated ICBs, along with the untreated PBS control, were injected peritumorally (intradermally beside the tumor). The study groups were the following: PBS, Abs only (∝-CTLA4 and ∝-PDL1 Abs), or Abs-cationic pep (∝-CTLA4-cationic pep and ∝-PDL1-cationic pep). Mice were humanely sacrificed when either the overall volume of the tumor exceeds 500 mm^3^, the size of the tumor in one dimension (length) exceeds 10 mm, or the body condition of the mice deteriorated significantly.

## ACKNOWLEDGMENTS

We thank the University of Texas at Austin Proteomics Facility, which is part of the Center for Biomedical Research Support for helping us with the protein identification analysis using LC-MS.

## AUTHOR CONTRIBUTIONS

R.P.M. and D.G. conceptualized the overall study. R.P.M. designed and performed experiments to conjugate and characterize peptides to the ICBs and estimate their binding to different proteins using ELISA. Y.P. designed and performed experiments to evaluate the off-tissue, on-target toxic effects of the peptide-conjugated Abs. M.M.L. and R.P.M. designed and performed experiments to investigate the immune cell profiling in the tumor and tumor-draining lymph nodes. R.P.M. designed and performed the antitumor efficacy experiments. M.S., E.Y.M., and R.F.A. helped with the animal experiments. Y.P., M.M.L. and R.P.M. analyzed the experiments. R.P.M., Y.P., and D.G. wrote the manuscript.

