## Supplemental Information for "Electrostatic-driven Interactions Enhance Intratumoral Retention and Antitumor Efficacy of Immune Checkpoint Blockade Antibodies"

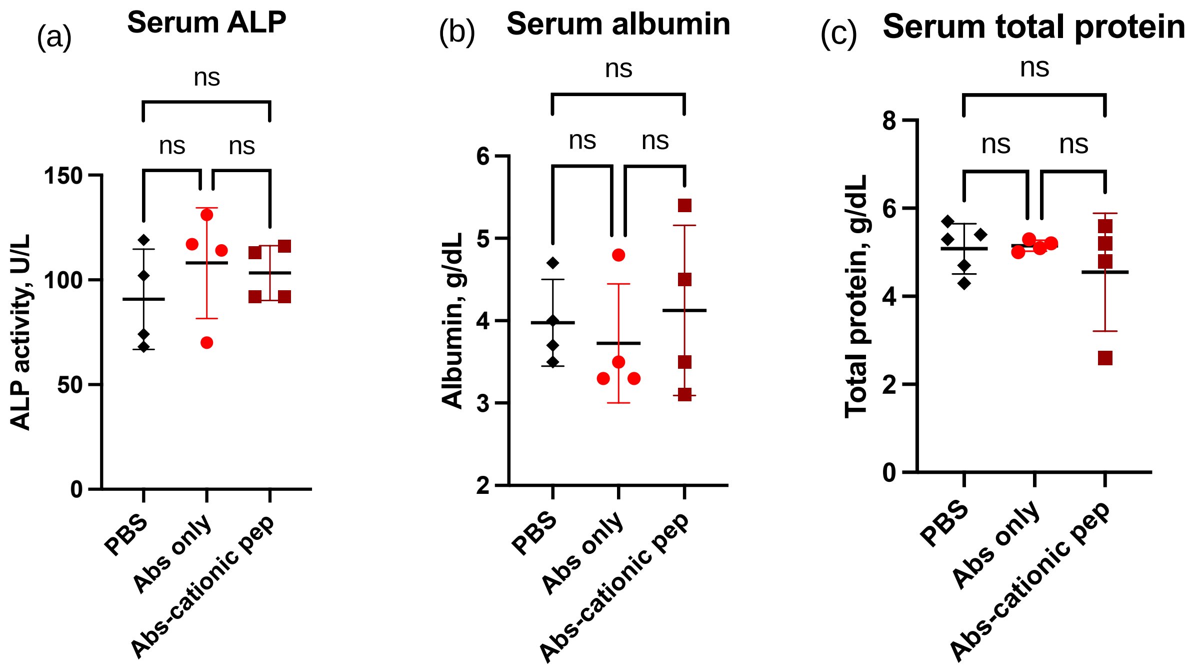


**Fig. S1.** No treatment-related hepatotoxicity was observed after a single administration of the cationic peptide-conjugated ICBs. Serum ALP activity, albumin, and total protein were measured

after blood collection on day 7. PBS p.t., n=5 for serum total protein; other treatment group, n=4; two experimental repeats.

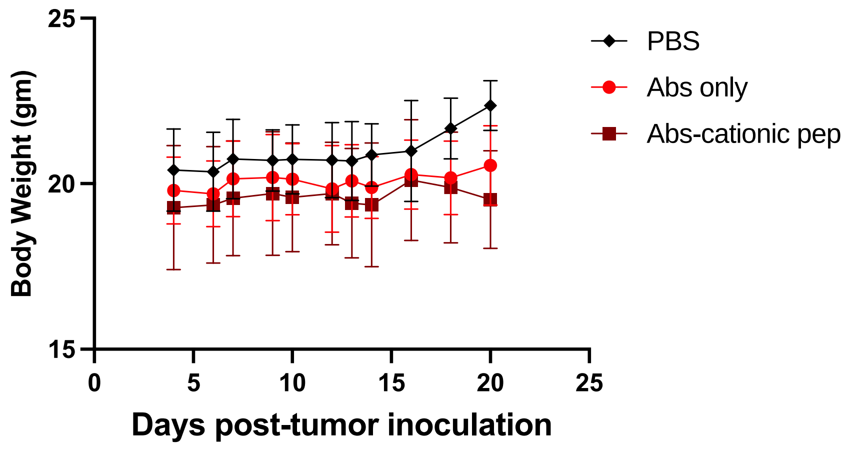


**Fig. S2.** The body weight of mice in each group was evaluated during the study duration. The average body weight of the mice was found to be maintained during the study.

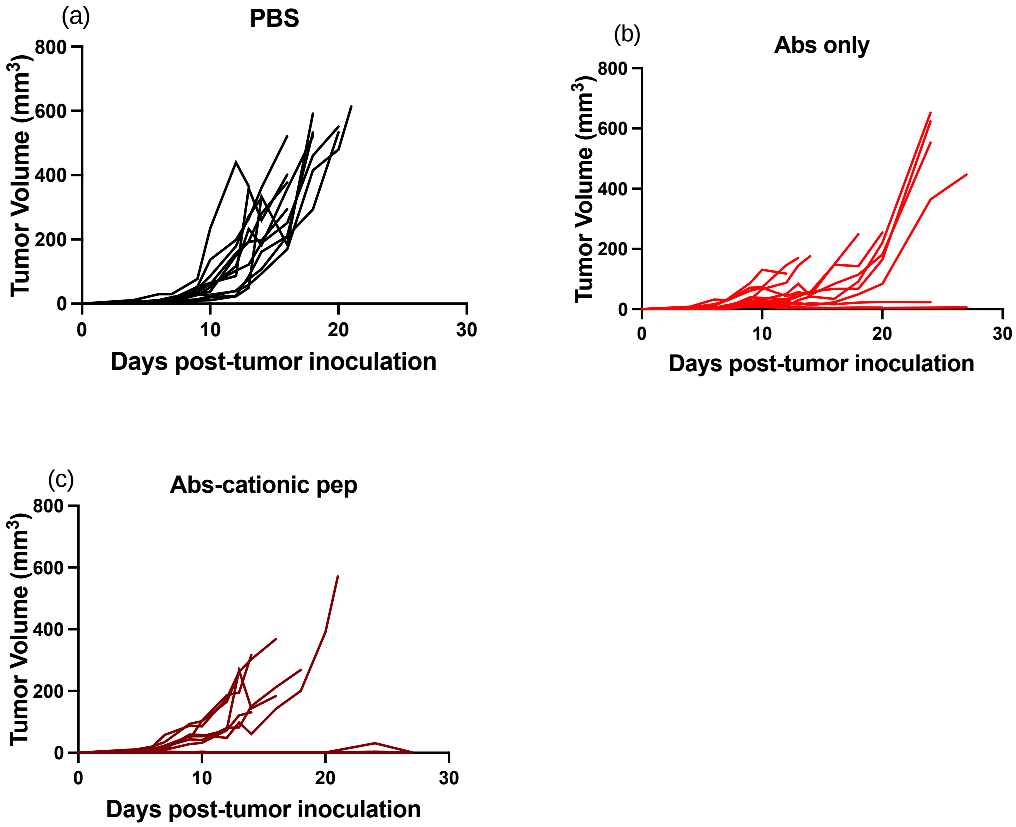


**Fig. S3.** Tumor regression of individual mice over the days of survival study for (a) PBS, (b) Abs only, and (c) Abs-neutral pep
